# Hippocampal cell- and circuit-specific differences in mitochondrial form and function

**DOI:** 10.64898/2025.12.16.694759

**Authors:** Mayd Alsalman, Lucy Turner, Katy Pannoni, Renesa Tarannum, Rutvi Desai, Sharon A. Swanger, Shannon Farris

## Abstract

Mitochondrial morphology varies by neuronal cell type and subcellular compartment; however, the functional significance of these differences is unclear. Hippocampal CA2 neurons are enriched for genes encoding mitochondrial proteins compared to CA1 neurons, suggesting a difference in metabolic demand across hippocampal circuits. However, whether CA2 neuron mitochondria are structurally or functionally distinct to support circuit-specific energy demands is unknown. Here we compared mitochondrial morphology, protein expression, and calcium levels across CA1 and CA2 circuits. We found mitochondria in CA2 dendrites were larger than mitochondria in CA1 dendrites. However, both subregions harbored larger mitochondria in the entorhinal cortex (EC)-contacting distal dendrites compared to CA3-contacting proximal dendrites. Together, these data demonstrate both cell type- and input-specific regulation of mitochondrial morphology that likely influences the function of these distinct circuits. To determine whether differences in mitochondrial fission or fusion account for cell and/or layer specific differences in morphology, we immunostained for OPA1 and MFF, which showed a general enrichment in distal dendrites relative to proximal dendrites, and an unexpected increase in CA1 distal dendrites compared to CA2 distal dendrites. To show whether these morphological differences result in functionally distinct mitochondria, we measured mitochondrial calcium levels in live slices. We found a striking enrichment of mitochondrial calcium levels in CA2 distal dendrites relative to proximal dendrites, and this layer-specific effect was significantly different from that in CA1 dendrites at baseline and after activity. Collectively, these findings reveal discrete morphological and functional differences in mitochondria across hippocampal subregions and dendritic layers, which likely confer unique circuit properties and/or vulnerabilities to disease.

## 1 NTRODUCTION

Mitochondria are energy-supplying units that are distributed throughout neuronal axons and dendrites to locally support metabolically demanding compartment-specific functions, such as synaptic transmission and synaptic plasticity. In mature neurons, mitochondria are the predominant and most efficient source of cellular energy, ATP, as well as other metabolites and lipids (Kann & Kovács, 2007). They also buffer cytosolic calcium, a critical mediator of synaptic signaling (Walters & Usachev, 2023). Thus, properly distributed mitochondria are highly important for proper synapse function (Duarte et al., 2023). Indeed, disruptions to the mitochondrial network are a primary driver of synapse dysfunction in a growing number of neurodevelopmental (Licznerski et al., 2020; Rojas-Charry et al., 2021; Shen et al., 2019; Tang et al., 2013; Xian et al., 2025), psychiatric (Altar et al., 2005; Flippo & Strack, 2017a; MacDonald et al., 2020; Pennington et al., 2008; Whitehurst & Howes, 2022), and neurodegenerative disorders (Chen et al., 2019; Ferree & Shirihai, 2012; Lee et al., 2022; Park et al., 2018; Pickett et al., 2018; Reeve et al., 2018; Wang et al., 2023).

Mitochondrial function is closely tied to its morphology, with fusion-induced changes resulting in higher mitochondrial ATP (Afzal et al., 2021; Seager et al., 2020; Suzuki et al., 2018) and buffering more calcium (Kowaltowski et al., 2019). Many studies have characterized heterogeneity in mitochondrial ultrastructure across cell types (Delgado et al., 2019; Faitg et al., 2021; Monzel et al., 2023) and compartments (Delgado et al., 2019; Donovan et al., 2024; Faitg et al., 2021; Lewis et al., 2018; Pannoni et al., 2023; Virga et al., 2024), however the functional implications of these differences remain unclear. Compared to neighboring hippocampal subregions, CA2 neurons are enriched for transcripts encoding mitochondrial proteins, including the mitochondrial calcium uniporter (MCU), suggesting differences in mitochondrial capabilities (Farris et al., 2019). Within CA2 neurons, there are dendritic layer-specific differences in mitochondrial morphology with distal dendrites in stratum lacunosum moleculare (SLM) harboring larger, more tubular mitochondria than mitochondria in proximal dendrites in stratum radiatum (SR) (Pannoni et al., 2023, 2025). These morphologically distinct populations of mitochondria precisely align with discrete inputs from CA3 onto SR dendrites and from entorhinal cortex (EC) onto SLM dendrites, suggesting circuit-specific differences in energy demand. A similar trend was reported in CA1 dendritic layers where mitochondria in CA3-contacting SR harbored smaller mitochondria than EC-contacting SLM (Lee et al., 2022) due to layer-specific differences in activity-driven mitochondrial dynamics (Virga et al., 2024). However, whether there are differences in mitochondrial morphology and function between CA2 and CA1 neurons is not known.

Given the enrichment of mitochondrial calcium signaling proteins in CA2 compared to CA1 (Farris et al., 2019; Pannoni et al., 2023, 2025), we hypothesized that CA2 dendrites harbor larger mitochondria that buffer more calcium than those in CA1 dendrites. Using adeno-associated viral vectors (AAVs) to label and monitor mitochondria, combined with cell- and dendritic layer-specific analyses, we found that mitochondria in CA2 dendrites are larger in area than mitochondria in CA1 dendrites, a difference that is not well explained by fission and fusion protein expression profiles. However, there was a general enrichment of fission and fusion factors in SLM of both subregions, which may reflect the increased mitochondrial content there compared to SR. With regards to mitochondrial function, we uncovered a significant enrichment of mitochondrial matrix calcium at baseline in CA2 SLM relative to SR that maps onto the larger, more abundant mitochondria with enriched MCU expression in CA2 SLM. CA1 dendrites had minimal layer differences in mitochondrial calcium levels. Collectively, these data suggest that synaptic inputs and cell-specific signaling drive differences in mitochondrial form and function across CA1 and CA2 dendrites, which may be important for circuit-specific properties supporting hippocampal dependent memory.

## 2 RESULTS

### CA2 neurons harbor larger mitochondria that occupy more of the dendrite than CA1 neurons

To investigate morphological differences in dendritic mitochondria within and across cell types, we sparsely labeled mitochondria in CA1 and CA2 neurons by infusing an adeno-associated viral vector (AAV) expressing cre (AAV9-hSyn1-cre) into the hippocampus of MitoTag mice (Fecher et al., 2019) crossed with a TdTomato reporter (Madisen et al., 2010). This enabled visualization of GFP-labeled mitochondria within TdTomato filled dendrites (Fig 1A-C). At the population level, individual mitochondria in secondary and tertiary dendrites of CA2 SLM were the largest in area compared to those in CA1 SLM, CA2 SR and CA1 SR (Fig. 1D). At the animal level, dendritic mitochondria in CA2 were larger than those in CA1 (paired two-way ANOVA, main effect of subregion F(1, 7) = 33.32, p=0.0007, N=8 mice) (Fig. 1 E). Consistent with previous studies (Pannoni et al., 2023; Virga et al., 2024), dendritic mitochondria in SLM were larger than those in SR in both subregions (paired two-way ANOVA, main effect of layer F(1, 7) = 86.18, p<0.0001) (Fig. 1 E). To exclude subregion differences based on cell size (Chevaleyre & Siegelbaum, 2010; Ishizuka et al., 1995) or plane of sectioning (Ishizuka et al., 1995), we compared segmented dendrite areas and found no significant differences between CA1 and CA2 (Fig. 1 F). To compare the percent area of the dendrite occupied by mitochondria, we divided the total area of all mitochondria within a dendritic segment by the area of that dendritic segment. The mitochondrial differences across CA1 and CA2 and between dendritic layers within each cell type were maintained when normalized by dendrite area, albeit slightly diminished (Fig. 1G; paired two-way ANOVA; main effects of subregion F(1, 7) = 16.11, p=0.0051, and layer F(1, 7) = 39.96, p=0.0004, N=8 mice). CA2 SLM had the highest % dendritic area occupied by mitochondria (45.47 ± 2.65%) followed by CA1 SLM (38.29 ± 2.61%), CA2 SR (35.81 ± 2.2%) and CA1 SR (30.79 ± 1.64%) (Fig. 1F, pairwise statistics reported on the plot). These data indicate that mitochondria size differs across hippocampal inputs and cell types.

**Fig. 1:**
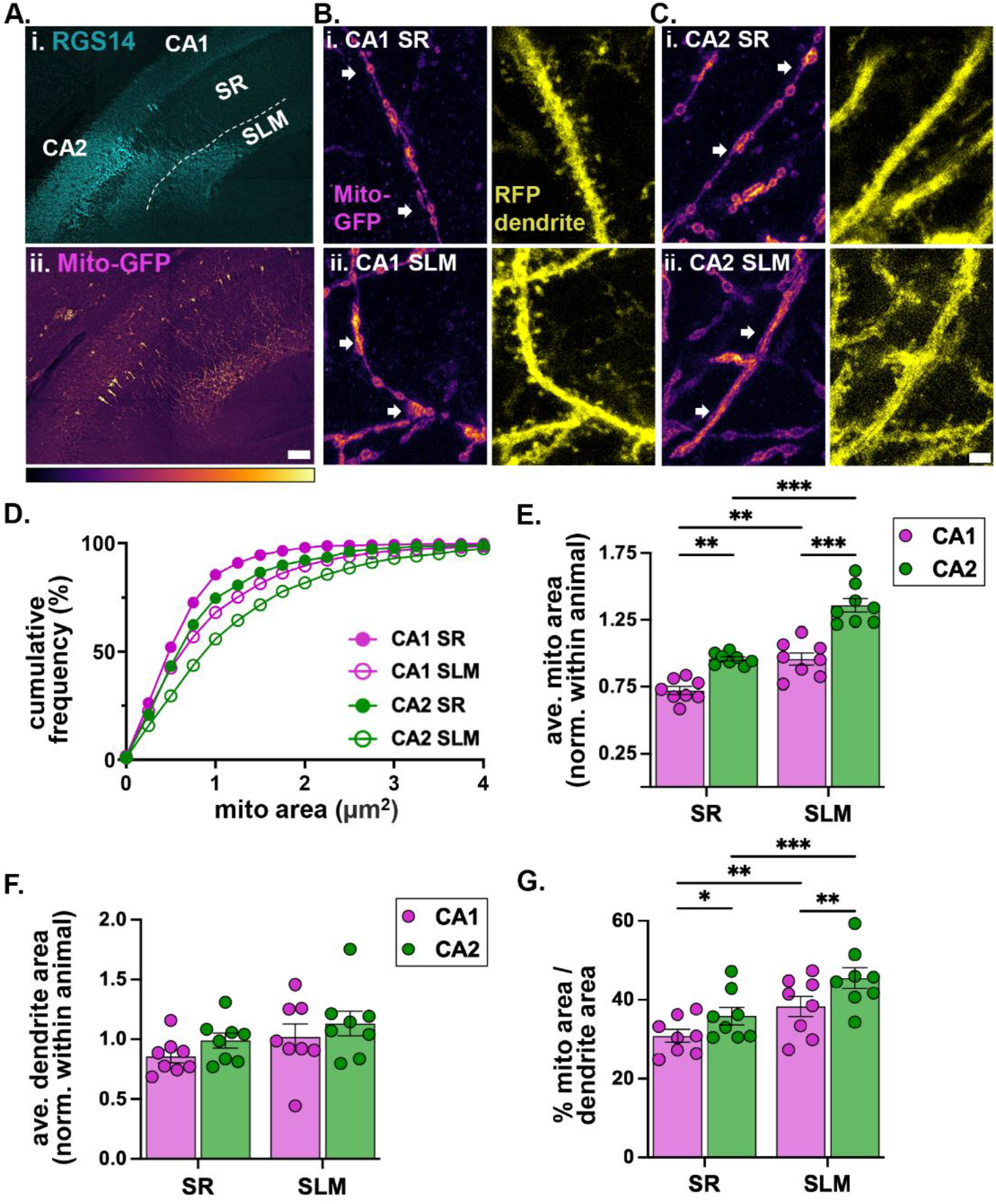
Dendritic mitochondria are larger in CA2 than in CA1 and occupy a greater percent of the dendrite. A.Representative coronal image of the hippocampus showing CA2 neurons (labeled with RGS14) and sparse, cre-dependent labeling of mitochondria (Mito-GFP) B.Representative high magnification images of Mito-GFP and RFP-labeled secondary dendrites in CA1 SR (i) and SLM (ii). C.Representative high magnification images of Mito-GFP and RFP-labeled secondary dendrites in CA2 SR (i) and SLM (ii). MitoTag-GFP images are scaled to the same fluorescence range. RFP images are scaled to visualization. D.Histogram showing the percent cumulative frequency of non-normalized mitochondria areas in CA1 and CA2 SR and SLM. CA1 SR n = 1866 mitochondria from 111 dendrites, SLM n = 2387 mitochondria from 122 dendrites; CA2 SR n = 1653 mitochondria from 96 dendrites, CA2 SLM n = 2159 mitochondria from 119 dendrites from 8 mice. E.Bar plot of average mitochondrial areas per animal (N= 8 mice). Data are normalized to each animal average. (paired two-way ANOVA, main effect of subregion F(1, 7) = 33.32, p=0.0007 and main effect of layer F(1, 7) = 86.18, p<0.0001, Fisher’s LSD posthoc tests are shown on the plot) F.Bar plot of average dendrite areas per animal (N= 8 mice). Data are normalized to each animal average. (paired two-way ANOVA, no effect of subregion F (1, 7) = 4.069, p=0.0835, no effect of layer F(1, 7) = 0.9857 p=0.3539) G.Bar plot of percent area of the dendrite occupied by mitochondria. (paired two-way ANOVA, main effect of subregion F(1, 7) = 16.11, p=0.0051, and layer F(1, 7) = 39.96, p=0.0004, Fisher’s LSD posthoc tests are shown on the plot)*p <0.05; **p<0.01; ***p<0.001. Scale bars = 100 µm (A), 2 µm (C). Error bars are average ± SEM.

### Mitochondrial fission and fusion proteins are abundant in CA1 SLM

To investigate whether differences in mitochondrial areas in CA1 and CA2 are due to differences in fission and fusion, we immunostained for fusion protein, optic atrophy 1 (OPA1), and fission protein, mitochondrial fission factor (MFF). We compared OPA1 and MFF fluorescence intensity as a measure of protein abundance across subregions and dendritic layers (Fig. 2). Strikingly, OPA1 showed a prominent band in CA1 SLM (Fig. 2AB), which was contrary to our initial expectation that CA2 would be more fusion dominant based on increased mitochondrial areas compared to CA1 (Fig. 1). OPA1 intensity significantly differed by subregion and dendritic layer (F(1,7) =6.623, p=0.0368), layer (F(2,14) =52.02, p<0.0001), and showed an interaction effect of subregion x layer (F(2,14) =86.69, p<0.0001) (Fig. 2C). The highest OPA1 intensity was in CA1 SLM despite lower OPA1 intensity in CA1 cell bodies (stratum pyramidale, SP) compared to CA2 cell bodies (p<0.0001, Tukey’s posthoc test) (Fig. 2C). For both subregions, OPA1 intensity was higher in SLM compared to SR (p<0.0001) (Fig. 2C). In contrast, MFF intensity (Fig. 2DE) only significantly differed by layer (F(2, 14) = 306.9, p<0.0001) with an interaction effect of subregion x layer (F(2, 14) = 32.21, p<0.0001) (Fig. 2GH). The highest MFF intensity was in CA2 SP, which was significantly higher than CA1 SP (p<0.0001, Tukey’s posthoc test), yet paradoxically lower in CA2 SLM relative to CA1 SLM (p=0.0003).

**Fig. 2:**
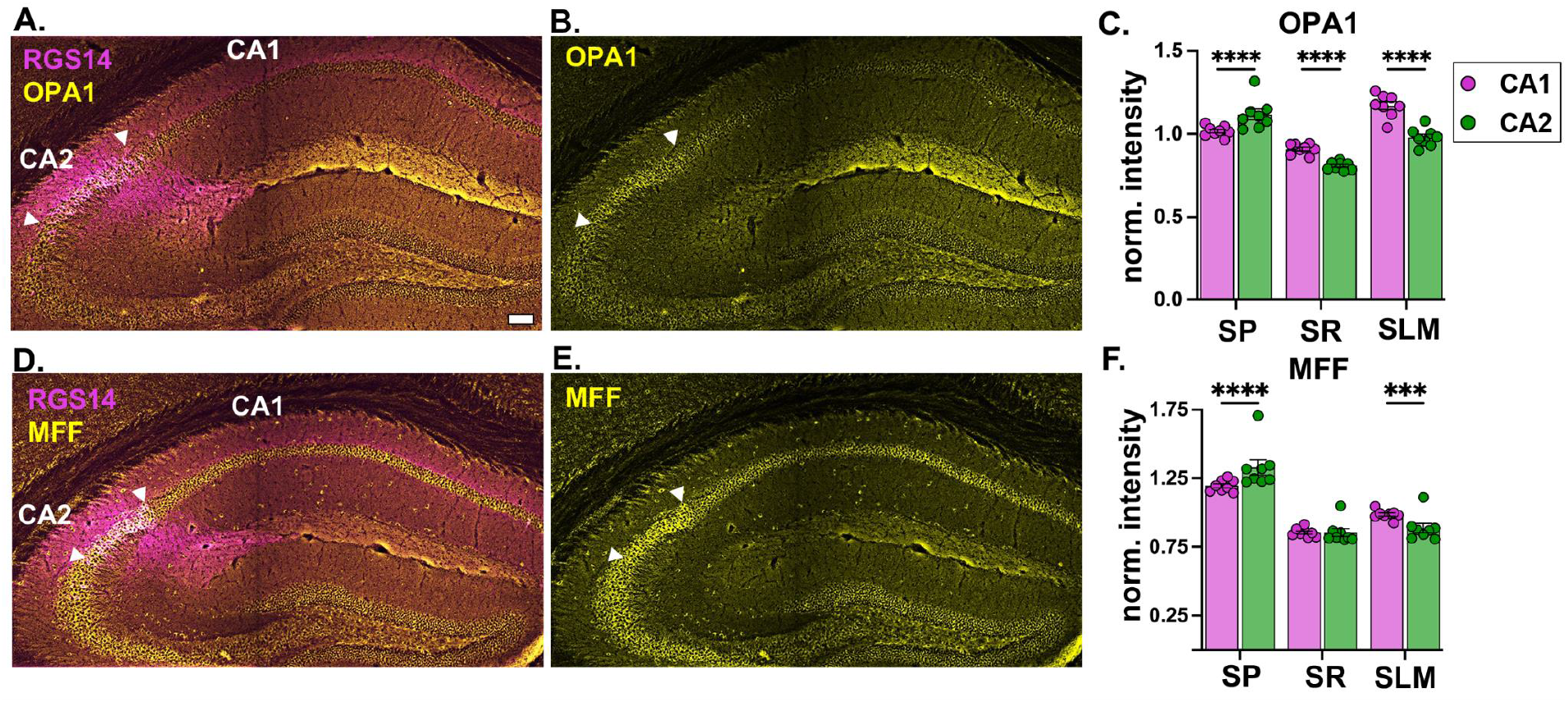
Mitochondrial fusion and fission proteins OPA1 and MFF are enriched in CA1 SLM. A.Representative coronal tile image of OPA1 and CA2 marker RGS14 in the hippocampus. Arrowheads denote CA2 cell body borders. B.The same image as in A with only OPA1. C.Bar plot of OPA1 fluorescence intensity in CA1 and CA2 SP, SR and SLM normalized to the animal average. (Paired two-way ANOVA, main effect of subregion (F(1,7) =6.623, p=0.0368), layer (F(2,14)=52.02, p<0.0001), and interaction effect of subregion x layer (F(2,14) =86.69, p<0.0001, N = 8 mice, Tukey’s posthoc tests are shown on the plot). D.Representative coronal tile image of MFF and CA2 marker RGS14 in the hippocampus. Arrowheads denote CA2 cell body borders. E.The same image as in D with only MFF. F.Quantification of MFF fluorescence intensity in CA1 and CA2 SP, SR and SLM normalized to the animal average. (Paired two-way ANOVA, main effect of subregion (F(2, 14) = 306.9, p<0.0001) with an interaction effect of subregion x layer (F(2, 14) = 32.21, p<0.0001, N = 8 mice, Tukey’s posthoc tests are shown on the plot). ***p<0.001 ; ****p<0.0001. Scale bar = 100 µm. Error bars are average ± SEM.

### Hippocampal subregion- and layer-specific differences in dendritic mitochondrial calcium signaling

In order to test whether mitochondria in CA1 and CA2 dendrites are functionally different, we measured mitochondrial calcium levels using a mitochondrial matrix targeted calcium indicator, mito-RGECO, in live slices (Fig, 3). To compare dendritic mitochondrial calcium signals across neighboring subregions, we unilaterally infused AAVs into CA2 or CA1 in separate cohorts of mice to prevent contaminating signals from upstream or commissural axons within the neuropil. To assess whether differences in cytosolic calcium levels contribute to differences in mitochondrial calcium levels, we co-infused the cytosolic calcium sensor, cyto-GCaMP6f. CA2 neurons have a unique tropism for transducing AAVs (Alexander et al., 2024) causing differences in AAV expression across subregions. To account for this, we extended the AAV incubation timeline for CA1 transduction (see methods) to ensure similar and sufficient labeling per subregion and statistically compared layer-specific differences between CA1 and CA2 using within cell type ratios. Moreover, due to anatomical differences in the plane of dendrites in CA1 (dorsoventral) and CA2 (septotemporal), we modified the plane of sectioning accordingly (CA1 coronal, CA2 horizontal). This resulted in fully in-plane transduced neurons in at least one and at most three slices (300 micron thick) per mouse. At baseline, there was a slight elevation of dendritic mitochondrial calcium signals in CA1 SLM relative to CA1 SR (Fig. 3Ai, B). In contrast, dendritic mitochondrial calcium signals in CA2 SLM were obviously higher than in CA2 SR (Fig. 3Ci, D) and this layer-specific effect was statistically different than in CA1 dendrites (two-way ANOVA, main effect of subregion, F (1,11) = 8.612, p=0.0136, baseline CA1 vs CA2, p= 0.0193, Fishers LSD, Fig. 3F). Upon depolarization with bath applied KCl, mitochondrial calcium signals increased throughout CA1 and CA2 dendrites (Fig. 3Aii, Cii, B-E), such that the subregion- and layer-specific differences remained similar to baseline (no main effect of treatment, p=0.6695; no interaction effect of treatment x subregion, p =0.1014, Fig. 3F). No differences were observed in cytosolic calcium levels that would explain the layer-specific differences in mitochondrial calcium levels (Fig. 3G-J).

**Fig. 3:**
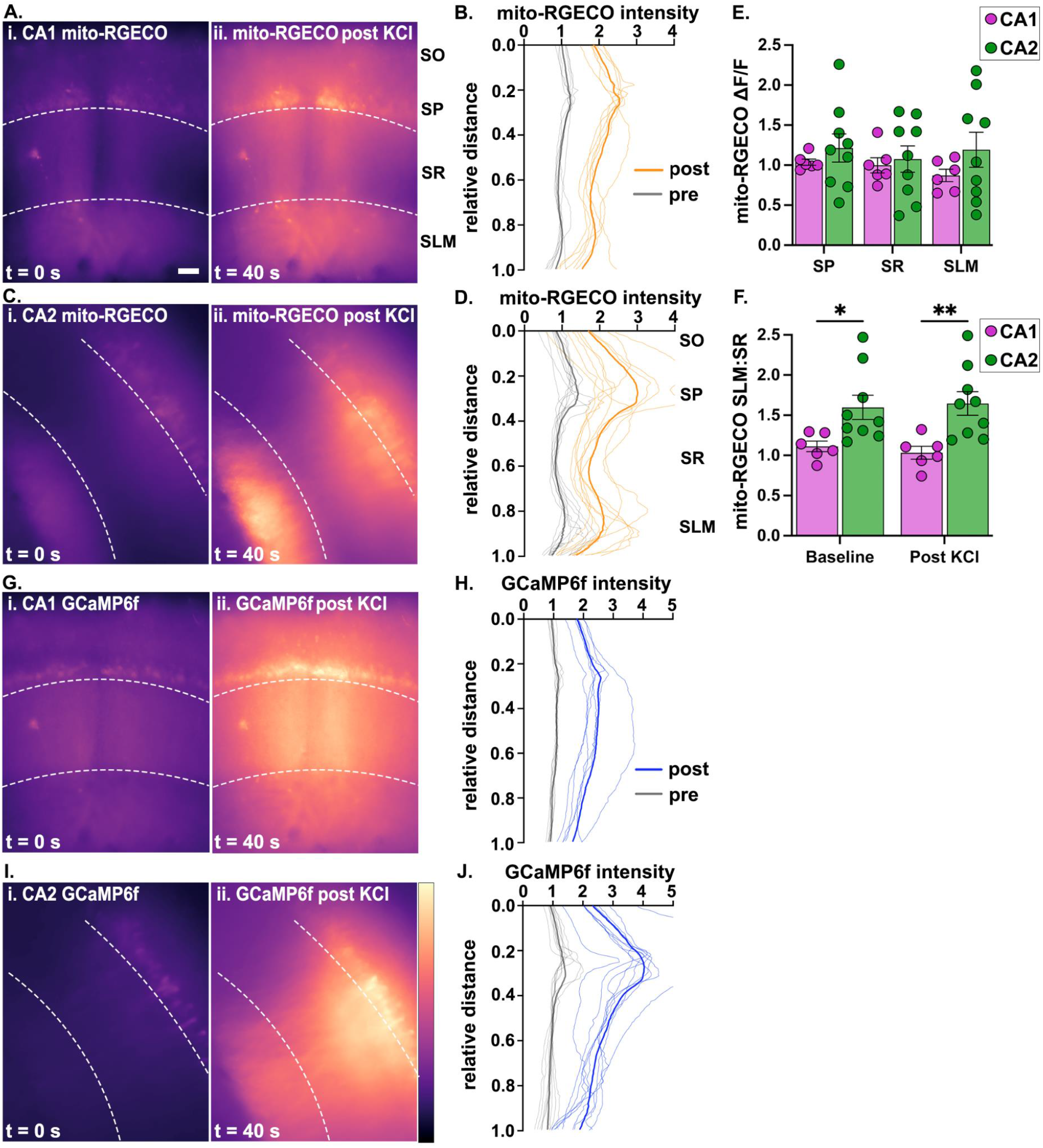
Mitochondrial calcium is enriched in CA2 SLM of live slices. A.Representative coronal live slice images of CA1 mitochondrial calcium pre (i) and post KCl treatment (ii). B.Normalized average mito-RGECO intensity line plots in CA1 layers relative to distance from stratum oriens (SO, 0.0). The thin lines represent the average trace per mouse normalized by the average intensity within each slice. Multiple slices from the same mouse were averaged to produce a single line per mouse. The thicker line is the average of all animals. (N=6 mice, 11 slices). C.Representative horizontal live slice images of CA2 mitochondrial calcium at baseline (i) and post KCl treatment (ii). D.Normalized average mito-RGECO intensity line plots in CA2 layers as in B. (N=9 mice, 12 slices). E.Quantification of KCl-induced change in mitochondrial calcium fluorescence normalized to baseline (ΔF/F) plotted as an average per animal (N=6-9 mice, 1-3 slices per mouse, 11 CA1 slices, 12 CA2 slices). No significant differences in subregion, layer, or interactions (two-way ANOVA). F.The ratio of mitochondrial calcium signals in SLM to SR at baseline and post KCl treatment (two-way ANOVA, main effect of subregion, F (1,13) = 8.244, p=0.0136, Fisher’s LSD posthoc tests are shown on the plot). G.Representative coronal live slice images of CA1 GCAMP6f cytosolic calcium levels pre (i) and post KCl (ii). The same slice is shown as in A. H.Normalized average GCaMP6f intensity line plots in CA1 layers as in B. I.Representative horizontal live slice images of CA2 GCAMP6f cytosolic calcium levels pre (i) and post KCl (ii). The same slice is shown as in C. J.Normalized average GCAMP6f intensity line plots in CA2 layers as in B. Note the lack of a peak in SLM relative to SR as seen in D. *p<0.05 p**<0.01. Scale = 100 µm. Error bars are average ± SEM. The brightness of images in A, C, G & I are optimized per slice.

### Anatomical differences contribute to enriched mito-RGECO signal in CA2 SLM

To investigate whether the functional enrichment of mitochondrial calcium in SLM relative to SR in live slices is driven, in part, by anatomical differences across subregions or dendritic layers, we perfused mice infused with AAV-mito-RGECO as above and immunolabeled the mito-RGECO signal with an RFP antibody (Fig. 4A). Live imaging of mito-RGECO signals reflects the relative amount of calcium within the mitochondrial matrix, whereas immunolabeling for mito-RGECO signals reveals the total amount of mito-RGECO protein expressed within the mitochondrial matrix. Thus, the immunolabeled mito-RGECO signal can be used to quantify layer differences in mitochondrial distribution within a subregion. The immunolabeled mito-RGECO signal was restricted to the mitochondrial matrix as evidenced by surrounding outer mitochondrial membrane tagged GFP (Fig 4B). The CA1/CA2 border was targeted for AAV infusion to quantify mito-RGECO fluorescence in both subregions in the same mice. To control for differences in transduction efficiency (see above), we normalized fluorescence intensities in SR and SLM within the same subregion (Fig. 4CD). To determine how mitochondrial distribution varies by dendritic layer, line plots were used to measure the mito-RGECO fluorescence intensity across CA1 and CA2, beginning in stratum oriens (SO) through SLM (Fig. 4D). In order to compare layer differences across subregions, we plotted the SLM:SR ratios (Fig. 4E). In CA2, the mito-RGECO signal was enriched in SLM compared to SR (ave. SLM:SR ratio ± SEM, 1.62 ± 0.09), whereas there was no difference between SLM and SR in CA1 (0.91 ± 0.07) (Fig 4E). This layer-specific difference in CA2 was statistically higher than observed in CA1 (unpaired two-tailed t test, effect of subregion, p=0.0001), suggesting asymmetric mitochondrial distribution plays a role in the enrichment of mitochondrial calcium signals in SLM of CA2, but not in CA1.

**Fig. 4:**
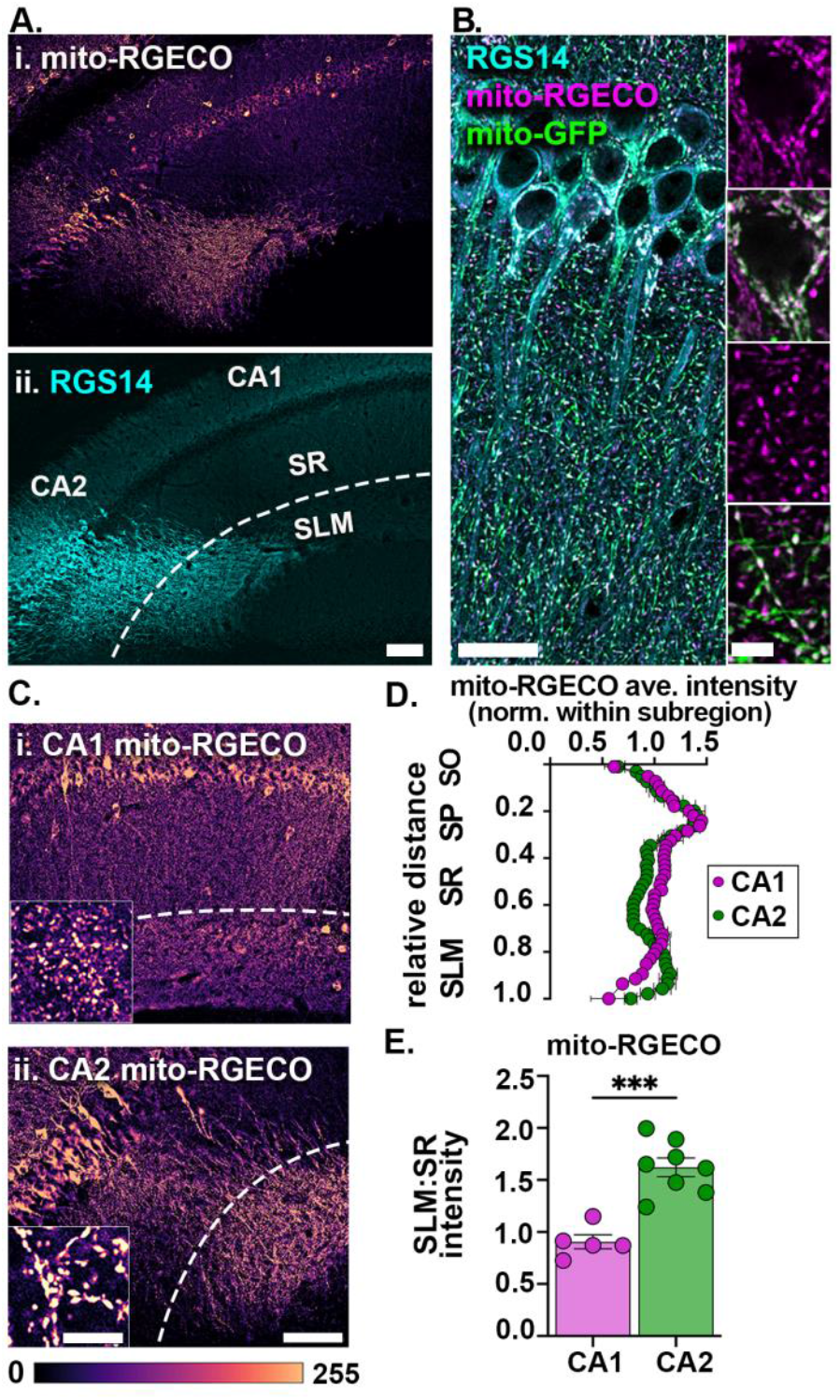
mito-RGECO expression is enriched in CA2 SLM in fixed sections. A.Representative tile image of mito-RGECO (i) and CA2 marker RGS14 (ii) in a perfusion fixed, coronal section of the hippocampus (dashed line denotes the border of SR and SLM). B.Representative high magnification image of RGS14 labeled CA2 neurons (cyan) expressing mito-RGECO (magenta) and sparse, cre-dependent labeling of MitoTag-GFP (green). Insets of a CA2 cell soma and proximal dendrites. Note the mito-RGECO signal is localized to the mitochondrial matrix surrounded by the GFP-labeled outer membrane. C.Representative images of mito-RGECO in CA1 (i) and CA2 (ii). Insets show mito-RGECO in distal dendrites. D.Normalized average mito-RGECO intensity line plots across CA1 and CA2 layers relative to distance from stratum oriens (SO). Data points are normalized by the average intensity within a subregion per animal. Thus, CA1 and CA2 data should not be directly compared, but rather the layer differences within each subregion. The data shown are animal averages ± SEM. (CA1 N=5 mice, CA2 N=7 mice). E.Average ratio of SR to SLM mito-RGECO fluorescence intensities. (unpaired two tailed t-test, CA1 N=5 mice, CA2 N=7 mice) ***p<0.0001. Scale bars = 100 µm (A, C), 20 µm (B), 5 µm (B, C insets). Error bars are average ± SEM.

We also considered whether differences in dendritic arborization within SLM could be contributing to the enriched mitochondrial calcium signaling (Sup. Fig 1). Due to orientation, some slices have more or less labeled CA2 dendrites in SLM (Sup. Fig 1B). Regardless, the net average signal of mito-RGECO exceeded any effect of the dendritic plane. We confirmed the mito-RGECO enrichment in CA2 SLM in coronal live slices as well (N= 4 slices from 2 mice, data not shown). In CA1, sparse hSyn1-driven expression of TdTomato resulted in an apparent band of signal in SLM (Sup. Fig 1CD). While some of the signal appeared to be from CA1 dendrites, especially near the infusion site (Sup. Fig 1D), the signal spread several hundred micrometers beyond the epicenter, suggesting the source was heavily ramified in SLM. We did not detect any extrahippocampal afferent labeling, e.g. nucleus reuniens of the thalamus (Wouterlood et al., 1990), that would suggest transynaptic labeling contributes to the banding. One possibility is the OLM interneurons, which are positioned in SO and send axonal projections that ramify in SLM (Thulin et al., 2025). In our study, a small subset of nonpyramidal neurons in SO are transduced with mito-RGECO (Sup. Fig 1AB). Additional studies using methods selective for excitatory neuron mito-RGECO are needed to determine the extent of their contribution to mitochondrial calcium signals in SLM.

**Sup. Fig. 1:**
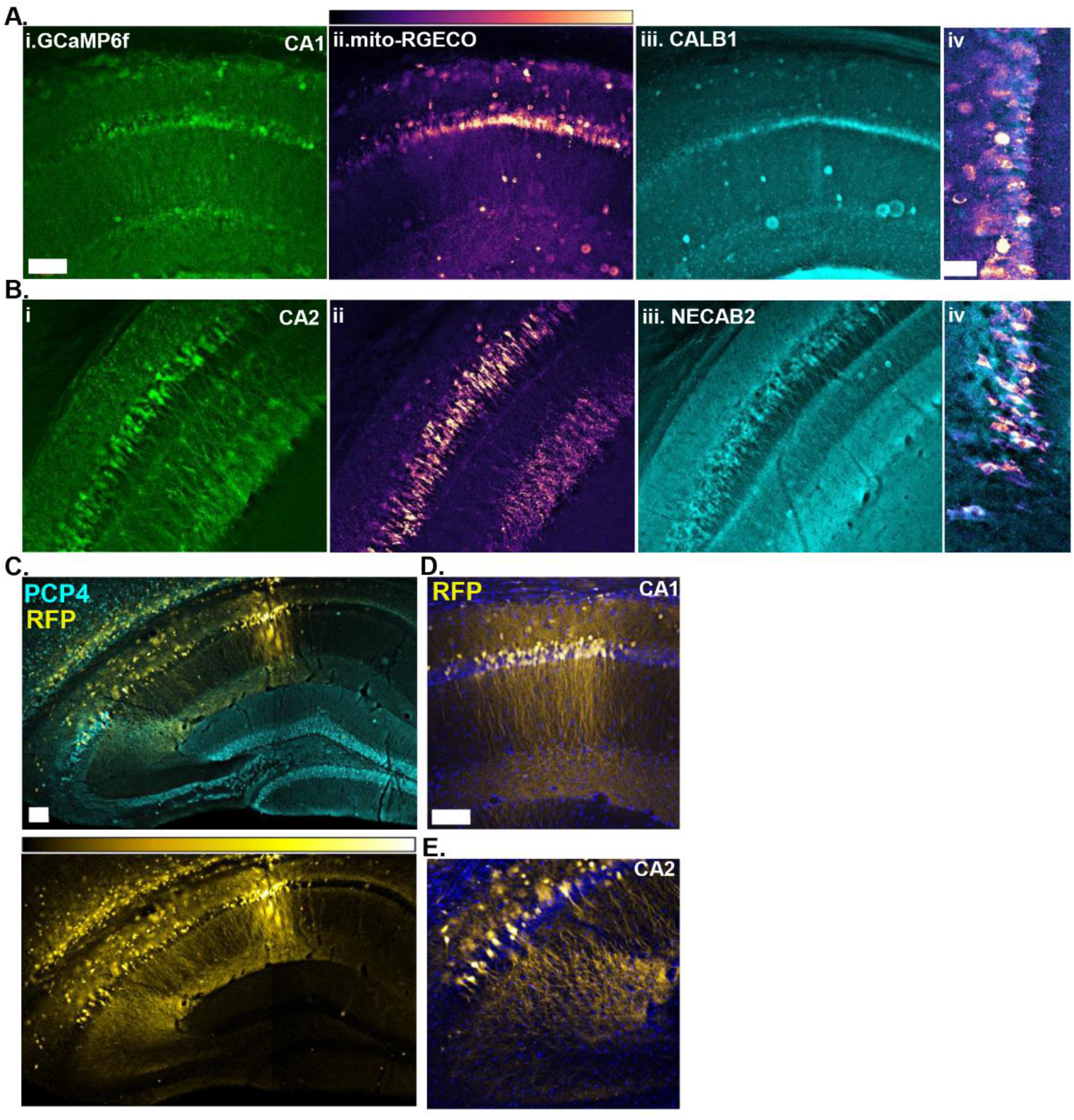
Validation of hSyn1-driven CA1 and CA2 mito-RGECO and GCaMP6f labeling. A.Post-fixed CA1 coronal slice stained for GFP (GCaMP6f, i), RFP (mito-RGECO, ii), and CA1 marker CALB1 (iii). Zoomed inset of SP (iv) to show mito-RGECO expression in CALB1 expressing CA1 neurons. B.Post-fixed CA2 horizontal slice stained for GFP (GCaMP6f, i), RFP (mito-RGECO, ii), and CA2 marker NECAB2 (iii). Zoomed inset of SP (iv) to show mito-RGECO expression in NECAB2 expressing CA2 neurons. C.Representative tile image of sparse hSyn1-cre driven tdtomato labeling of CA1 and CA2 neurons in a coronal section. CA2 neurons are labeled with PCP4. D.Higher magnification of CA1 to show dendritic banding in SLM. Nuclei are labeled with DAPI. E.Higher magnification of CA2 to show dendrites are out of plane. Nuclei are labeled with DAPI. Scale bars = (Ai) 100 µm, (Aiv) 50 µm, (C) 100 µm, (D) 100 µm

## 3 DISCUSSION

In this study, we investigated mitochondrial differences across hippocampal circuits, focusing on subregions CA1 and CA2. Using viral vector-based approaches combined with light microscopy, we visualized mitochondria, fission and fusion proteins, and mitochondrial matrix calcium levels in proximal (SR) versus distal (SLM) dendrites, which represent distinct circuits, receiving inputs from CA3 and EC, respectively. We found that mitochondria in CA2 are larger and occupy a greater area within the dendrite compared to CA1. Within each of the subregions, mitochondrial area was increased in SLM compared to SR. We further found increased expression of fission factor MFF and fusion factor OPA1 in SLM, the latter of which showed significant banding in CA1 SLM, which may reflect cell- and layer-specific differences in mitochondrial dynamics. Lastly, we assessed mitochondrial function by measuring mitochondrial matrix calcium in live slices. At baseline, there was a striking band of enriched mitochondrial matrix calcium in CA2 SLM and a less prominent band in CA1 SLM. When analyzing fixed tissues, some of this banding could be attributable to anatomical differences, including increased mitochondrial content in CA2 SLM and dendritic arborization in SLM of both subregions.

Activity-induced changes in cytosolic and mitochondrial calcium signals corroborated the baseline enrichment of mitochondrial calcium in CA2 SLM. Taken together, our data indicate that cell intrinsic and input-specific mechanisms contribute to the morphological and functional differences in dendritic mitochondria across hippocampal circuits. Dendritic mitochondrial heterogeneity may support the diversity of synaptic functions and connectivity patterns in different subregions supporting hippocampal-dependent memory. It may also contribute to differential circuit susceptibility to brain disorders.

### Dendritic mitochondria in CA2 are larger than in CA1

Mitochondrial area and dendritic occupancy were higher in CA2 compared to CA1, supporting the notion that CA2 mitochondria may be functionally distinct from those in CA1. However, comparing across cell types requires equivalent reporter expression. CA2 neurons have a unique tropism for most AAV serotypes due to enriched expression of the AAV receptor (AAVR; Alexander et al., 2024). This could, in theory, result in higher expression of the floxed stop mito-GFP due to earlier, more efficient transduction, influencing apparent mitochondrial size. However, this should also apply to the tdTomato reporter used as the dendritic fill. Thus, normalizing the mitochondria area to the dendrite area addresses this point. Further, the reporter cell body intensities appeared similar across cell types suggesting against reporter expression differences.

Within each subregion, distal dendrites (SLM) harbored larger mitochondria that took up a greater percentage of dendrite compared to proximal dendrites (SR), consistent with previous findings in CA1 (Lee et al., 2022; Virga et al., 2024) and CA2 (Pannoni et al., 2023, 2025). An increase in mitochondrial density in distal dendrites relative to those more proximal to the soma has also been described in other neuronal systems (Donovan et al., 2024). However, our use of an outer mitochondrial membrane (OMM) label may inflate the apparent size of the mitochondria and/or make it difficult to resolve two closely opposed mitochondria with diffraction limited light microscopy. Thus, individual mitochondria areas may be overestimated, especially for the more densely packed mitochondria in distal dendrites. However, the relative differences across layers and cell types should be maintained as similar results were shown for CA1 using both matrix and outer membrane targeted labels (Virga et al., 2024).

Subregion differences in mitochondrial size may be driven by differences in cell size or dendritic anatomy. CA2 cell somas are larger than CA1 cell somas (Ishizuka et al., 1995) and have different dendritic lengths and branching patterns (Helton et al., 2019; Piskorowski & Chevaleyre, 2011). There is evidence that dendritic architecture affects mitochondrial distribution. Horizontal system neurons in the Drosophila visual system showed higher mitochondrial motility into larger dendritic branches compared to smaller branches (Donovan et al., 2024), suggesting a link between dendritic thickness and mitochondrial content. While we did not systematically compare dendritic areas in CA1 versus CA2, it is possible that CA2 dendrites are larger than those in CA1 resulting in larger mitochondria.

Dendritic branching patterns contribute to the integration of synaptic inputs (Spruston, 2008; Sun et al., 2014) which may also influence dendritic mitochondrial morphology. CA1 and CA2 receive the same synaptic input from CA3 onto SR and similar input from EC layers II/III onto SLM, but they differ in their postsynaptic responses (Chevaleyre & Siegelbaum, 2010; Sun et al., 2014, 2017, 2021; Zhao et al., 2007). Compared to CA1 and CA3, CA2 has the largest dendritic area dedicated to EC input (Sun et al., 2017) and the largest synaptic response to EC stimulation (Chevaleyre & Siegelbaum, 2010; Sun et al., 2014, 2017). Synapses from CA3 onto CA1 are well-known for their ability to readily express long-term potentiation (LTP), but those same synapses onto CA2 are resistant to LTP (Zhao et al., 2007). However, CA3-CA2 synapses have the capacity to undergo potentiation under certain conditions (Carstens & Dudek, 2019) indicating that plasticity at this synapse is highly regulated rather than absent (Dudek et al., 2016). Cell and layer-specific differences in mitochondria could conceivably contribute to plasticity differences across circuits. For example, mitochondria in CA2 distal dendrites are enriched for MCU (Pannoni et al., 2023) and MCU deletion specifically in CA2 neurons resulted in diminished LTP at CA2 distal synapses (Pannoni et al., 2025). However, MCU may be similarly required for LTP at CA1 synapses, but this has yet to be tested. Further studies are needed to identify whether mitochondrial heterogeneity influences circuit-specific LTP profiles.

### CA1 SLM expresses more OPA1 than CA2 SLM

Mitochondria morphology is regulated by fission and fusion dynamics that respond to cellular activity to meet metabolic demands (Seager et al., 2020). Both fission and fusion maintain mitochondrial integrity by getting rid of damaged mitochondria or combining the healthy parts of two mitochondria (Seager et al., 2020), influencing their shape (Itoh et al., 2019; Joaquim et al., 2025). Fusion is generally thought to result in larger mitochondria and can thus potentially lead to enhanced energy production through larger cristae surface area for electron transport chain complex assembly (Afzal et al., 2021; Seager et al., 2020). Here we uncovered a striking banding in fusion factor OPA1 in CA1 SLM compared to CA2 SLM. This was contrary to our expectation given that CA2 SLM harbors larger mitochondria than those in CA1 SLM. The increase in OPA1 does not appear to be solely explained by increased mitochondrial content, as seen with other mitochondrial markers like COX4 (Pannoni et al., 2023), as the band is less striking in CA2 SLM, which has higher mitochondrial content than CA1 SLM. Enriched OPA1 in SLM is consistent with it being a fusion dominant layer to explain its increase in mitochondrial mass (Virga et al., 2024), although the effect was relatively modest in CA2 SLM despite robust expression in CA2 SP. The cell- and layer-specific expression of MFF and OPA1, together with the mitochondrial morphology findings, suggest that mitochondrial dynamics is shaped by both intrinsic cellular properties as well as synaptic inputs.

While these findings are striking by eye, it is important to note that the immunofluorescence techniques used here are not sensitive enough to differentiate signals coming from dendrites versus traversing axons. These proteins are also present in axonal mitochondria (Ju et al., 2005; Lewis et al., 2018; Seager et al., 2020) and thus it is conceivable that some of the OPA1 enrichment may be due to expression in ECIII axons. When considering the significantly higher OPA1 levels in CA2 SP over CA1 SP, it is otherwise unclear why OPA1 would not be present at higher levels in CA2 dendrites. Looking at other fission and fusion proteins (such as DRP1 and MFN1/2, respectively) could provide more information about the mitochondrial dynamics in these circuits. Further, OPA1 and MFF immunostaining provide a snapshot of total protein abundance, and it is possible that their enzymatic activities also differ by subregion and/or layer. Staining for the active forms of these proteins versus the total levels may be more informative of their activity. Moreover, the presence of these proteins could be necessary for functions other than mitochondrial dynamics (Itoh et al., 2019; Joaquim et al., 2025), and even when one or all the components are present, the process may not take place; for example, even assembly of the entire fission machinery does not always lead to mitochondrial fission (Flippo & Strack, 2017b). Thus, the mere enrichment of OPA1 may not indicate that CA1 SLM is more fusion-dominant than CA2 SLM. Further studies are needed to clarify how differences in mitochondrial dynamics contribute to hippocampal cell- and circuit-specific differences in mitochondrial morphology.

### CA2 SLM has higher basal mitochondrial calcium than SR

Due to differences in AAV expression across cell types, we did not directly compare mitochondrial calcium levels across subregions. Instead, we compared layer-specific mitochondrial calcium levels within cell type by generating a ratio of SLM to SR at baseline and post-KCl peak intensities. There was a significantly higher SLM:SR ratio in CA2 than CA1, suggesting that mitochondria in CA2 distal dendrites have higher baseline levels of mitochondrial calcium relative to proximal dendrites than in CA1. Further, the minutes-long increase in mitochondrial calcium uptake in response to depolarization maintained this effect. This layer-specific profile was not explained by differences in cytosolic calcium as the baseline and post-KCL cytosolic calcium levels were uniform across dendritic layers. Because the mitochondrial calcium differences were detected at baseline, it is unlikely that the KCl-induced increase in cytosolic calcium (that occurs over the entire slice) contributed to these layer-specific differences. However, it is possible that the effects obtained with bath applied KCl masks or amplifies calcium signals in non-physiological ways. Additional studies are needed to uncover whether input-specific and/or synaptically evoked changes in cytosolic calcium similarly result in layer-specific differences in mitochondrial calcium.

The CA subregions have different cytosolic calcium handling properties, which may contribute to the observed differences in mitochondrial calcium levels. For instance, CA2 neurons exhibit lower calcium transients in dendritic spines in response to action potential firing compared to CA1 and CA3 neurons, and have higher calcium extrusion and buffering capacities (Simons et al., 2009). Enhanced mitochondrial calcium buffering may be an additional layer of regulation in CA2 to limit intracellular calcium signaling. It is also conceivable that the robust cytosolic calcium buffering and extrusion mechanisms in CA2 (Simons et al., 2009) result in a more fusion dominant profile leading to larger sized mitochondria that buffer more calcium in CA2 compared to CA1. We did not directly compare cytosolic calcium levels between CA1 and CA2 due to the lack of a ratiometric tag to control for differences in AAV expression. We were also unable to quantitate layer-specific differences in cytosolic calcium levels due to the very low baseline GCaMP6f signal to noise at 10x magnification. Further experiments with faster imaging kinetics are needed to directly compare differences in cytosolic calcium levels between CA1 and CA2 dendrites to determine how they may contribute to differences in basal or evoked mitochondrial calcium levels.

We previously reported a significant enrichment of MCU in CA2 compared to CA1, which was especially striking in EC-contacting SLM dendrites (Pannoni et al., 2023, 2025) precisely where we see elevated baseline and evoked mitochondrial calcium levels. Mitochondrial calcium uptake, which in neurons is predominantly accomplished via MCU, is essential for the modulation of ATP production (Pivovarova & Andrews, 2010).

Increased intramitochondrial calcium levels stimulate the activity of some tricarboxylic acid (TCA) cycle enzymes (Feofilaktova et al., 2025), which create the electron carriers that are essential for the electron transport chain and ATP production. In acute cortical and hippocampal slices, MCU-mediated calcium uptake scaled with increased action potential firing (Groten & MacVicar, 2022), suggesting MCU may couple increased cytosolic calcium levels to increased mitochondrial metabolism. shRNA depletion of MCU in rat hippocampal cell culture resulted in a loss of spine stimulation-induced mitochondrial ATP production near dendritic spines (Ghosh et al., 2025) suggesting MCU signaling is required to meet energetic demands of nearby synapses. Given that CA2 SLM dendrites are enriched for MCU and mitochondrial calcium signals, and that MCU is required for LTP at CA2 SLM synapses (Pannoni et al., 2025), we speculate that MCU facilitates a circuit-specific ATP profile required for CA2 SLM synapse function.

### Greater mitochondrial content contributes to elevated mitochondrial calcium signals in CA2 SLM

It is possible that mitochondrial calcium levels may appear enriched in distal layers, at least in part, due to an increased mitochondrial content in CA2 SLM shown both as percent mitochondrial occupancy and increased individual mitochondrial area (Fig. 1). This was clearly seen with the increased signal for mito-RGECO in CA2 SLM in fixed tissue immunolabeling (Fig. 4). However, a similar effect was not detected in CA1 SLM in fixed tissues despite the small increase in matrix calcium in live slices. It is possible that the increase in matrix calcium in live slices in SLM of both subregions is due to larger mitochondria having the capacity to store more calcium than smaller mitochondria (Kowaltowski et al., 2019), but, because CA2 SLM has larger denser mitochondria than CA1 SLM, the effects are more pronounced there in fixed and live slices.

Our data provide a potential basis for cell-specific differences in mitochondrial form and function. We speculate that these differences may be established and/or maintained to support the unique metabolic demands of each circuit. Because mitochondria play an essential role in triggering calcium-mediated cell death (Pivovarova & Andrews, 2010), differences in mitochondrial calcium signaling could conceivably impart sensitivity or resilience to this form of cell death. Consistent with this, disruptions in mitochondrial calcium handling are associated with neurodegenerative disease pathology (Calì et al., 2012). Thus, the cell-specific differences we describe for dendritic mitochondrial calcium levels may influence circuit-specific susceptibility to such pathologies.

## 4 METHODS

### 4.2 Animals

Sexually naive adult male and female wildtype mice on a C57BL/6 background were used at the ages specified within the methods of each experiment. Animals were group-housed under a 12:12 light/dark cycle with access to food and water ad libitum. All procedures were approved by the Animal Care and Use Committee of Virginia Tech and were in accordance with the National Institutes of Health guidelines for care and use of animals.

Male and female homozygous Mitotag^fl/fl^ (Rosa26-CAG-LSL-GFP-OMM, Jax # 032675; Fecher et al., 2019) or Mitotag^fl/fl^ x TdTomato^fl/fl^ (Ai14(RCL-tdT)-D, Jax # 007908) mice were bred to generate the data in Fig. 1.Mitotag^fl/fl^, TdTomato^fl/fl^, Mitotag^fl/fl^ x TdTomato^fl/fl^, and Amigo2-EGFP (Tg(Amigo2-EGFP)LW244Gsat/Mmucd, MMRRC# [033018-UCD](https://www.mmrrc.org/catalog/sds.php?mmrrc_id=33018)) males and females were used to generate the data in Fig 2-4. All mouse strains used are considered wildtype animals.

### 4.2 AAV Infusion

Prior to the start of surgery, mice were weighed and acclimated to the room before receiving an intraperitoneal (IP) injection of ketamine/dexdomitor cocktail (ketamine 50-100 mg/kg; dexdomitor 0.375-0.5 mg/kg) at a 1:1 ratio for anesthesia. Mice were then placed on a heat pad and an eye lubricant was applied before the hair around the surgical area was trimmed. The mice are then placed on a stereotaxic apparatus with heat provided throughout the procedure. Prior to making the incision, the area was disinfected with 3 cycles of betadine and 70% ethanol. Once the incision was made, the skull was cleaned and unilateral or bilateral burr holes were drilled over the relevant hippocampal region (see below). AAVs (0.3-0.5µl) were infused with a glass pipette and a syringe pump at a rate of 50-100 nL/min. The experimenter waited 5 min before pulling out the glass pipette at a rate of 0.5 mm/min. The mouse was then removed from the stereotaxic apparatus and the incision closed using surgical glue. Mice were given an IP injection of antisedan (1-2 mg/kg) and a subcutaneous injection of extended release buprenorphine (3.25 mg/kg). Mice were then allowed to recover on a heating pad until they were ambulatory.

#### 4.2.1 Mitotag experiments

pAAV-hSyn-Cre-P2A-dTomato was a gift from Rylan Larsen (Addgene viral prep # 107738-AAV9; http://n2t.net/addgene:107738; RRID:Addgene_107738). AAV9-hsyn-cre-tdTomato (2.6^10^10^ GC/mL) was infused into the border of CA1/CA2 (coordinates from Bregma: -2.2 mm AP; +/-1.8 mm ML; -1.5 mm DV) bilaterally in Mitotag^fl/fl^ or unilaterally in Mitotag^fl/wt^;TdTomato^fl/wt^ mice to limit contralateral axonal labeling. Mice were infused at 1-5 months old and perfused 2-3 weeks after surgery.

#### 4.2.3 Genetically encoded calcium indicator experiments

pAAV.Syn.GCaMP6f.WPRE.SV40 was a gift from Douglas Kim & GENIE Project (Addgene viral prep # 100837-AAV9; RRID:Addgene_100837). AAV2/9-synapsin-mitoRGECO1.0 was purchased from the CNP Viral Vector Core at the CERVO Research Center contribution (RRID:SCR_016477). AAV2/9-synapsin-mitoRGECO1.0 (Neurophotonics, cat. #491-aav5, 3^10^11^ GC/mL) and AAV9-synapsin-GCaMP6f (8.5^10^11^ GC/mL; Chen 2014) were unilaterally co-infused into hippocampal CA1 (coordinates from Bregma: -2.3 mm AP; +1.5 mm ML, -1.5 mm DV) or CA2 (-2.2 mm AP; +2.2 mm ML, -1.65 mm DV) of wildtype mice at 4-9 weeks of age. Slices were prepared from these mice 2-3 weeks (CA2) or 4 weeks (CA1) post infusion.

Intermediate coordinates (-1.8 mm AP; +1.8 mm ML, -1.5 mm DV) and similar incubation periods (2-4 weeks post infusion) were used for fixed tissue mito-RGECO experiments.

### 4.3 Tissue processing and immunofluorescence

Mice were given an overdose of sodium pentobarbital (150 mg/kg) and perfused with 15-20 mL of 4% paraformaldehyde for 4 min. Mice infused with calcium indicators were perfused with 2 mM CaCl_2_ for 90 seconds prior to PFA. Brains were post-fixed in 4% paraformaldehyde for at least 24 hr before being sectioned coronally at 40µm on a Leica vibratome (VT1000S). All animals were stained using a direct immunofluorescence protocol, except for one that used a TSA boost protocol in the rabbit-anti-RFP channel only to boost dendritic signal. Direct fluorescence included washes in 1x phosphate buffered saline (1x PBS) and a 10 min permeabilization step in 0.03% Triton in 1x PBS (PBST). Sections were then blocked for a minimum of 1 hour in 5% Normal Goat Serum in PBST (5% NGS), followed by incubation in the primary antibody in the same diluent for 18-24 hours at room temperature. Negative control sections underwent the same steps in each run except for the primary antibody incubations, as described below.

#### MitoTag IHC

Sections used to generate figure 1 were immunostained with chicken-anti-GFP (1:2000, Abcam cat# 13970, RRID: AB_300798). Sections that came from a Mitotag^fl/fl^ mouse were additionally stained with rabbit-anti-RFP (1:500-1:2000, Rockland cat# 600-401-379, RRID: AB_2209751) to boost dendritic labeling. Guinea pig-anti-PCP4 (1:1000, Synaptic Systems cat# 480004, RRID: AB_2927387) was used as a CA2 maker. Sections then underwent PBST washes and a 30-minute block in 5% NGS then were incubated in secondary antibodies for 2 hours at room temperature, followed by PBST washes and a final wash in 1x PBS. Secondary antibodies included goat-anti-chicken (Alexa 488, 1:500, Invitrogen cat# A11039), goat-anti-rabbit (Alexa 546, 1:500, Invitrogen cat# A11035), and goat-anti-guinea-pig (Alexa 633, 1:500, Invitrogen cat# A21105). One mouse was with inadequate RFP/TdTomato signal was boosted using a Tyramide Signal Amplification (TSA) immunostaining protocol with a TSA Plus Cyanine 3 kit (Akoya Biosciences cat# NEL744001KT) in the RFP/TdTomato channel only. This protocol included washes in Tris Buffered Saline (TBS) and an optional 15-minute permeabilization in 0.03% triton in TBS (TBST), followed by a 1-hour block in TSA blocking buffer before incubating in primary antibody overnight at room temperature. Sections were then washed in TBST multiple times then the signal was quenched with 2% H2O2 in TBS for 10 minutes. After a second 15-minute block, sections were incubated in secondary antibodies at room temperature for 2 hours. In addition to goat-anti-chicken and goat-anti-guinea-pig-633 described above, goat-anti-rabbit-HRP (1:500, cat# 111-035-144) was added, followed by TBST washes. Sections were then incubated in TSA-Cy3 1:50 in PE 1x amplification diluent for 20 minutes. Both CA1 and CA2 dendrites were quantified from the TSA-boosted RFP-labeled sections, and the normalized data did not significantly differ from those without TSA. Sections were mounted and coverslipped with prolong gold fluorescence media with 4’,6-diamidino-2-phenylindole (DAPI; Invitrogen cat# P36931) and/or an additional 10-minute DAPI wash at 1:10,000 in 1x PBS prior to coverslipping.

#### Fission/Fusion IHC

Sections used to generate figure 2 were immunostained following the direct fluorescence protocol but included an antigen retrieval step done through boiling free-floating sections in 1mL of nanopure water prior to permeabilization. Sections were immunostained with mouse-anti-RGS14 (1:500, NeuroMab cat# 75-170, RRID: AB_2877352) as a CA2 marker as well as rabbit-anti-OPA1 (1:200, Proteintech cat# 27733-1-AP, RRID: AB_2810292) or rabbit-anti-MFF (1:500, Proteintech cat# 17090-1-AP, RRID: AB_2142463). The secondary antibodies included the following: goat-anti-mouse (Alexa 488, 1:500, Invitrogen cat# A11029) and goat-anti-rabbit (Alexa 546, 1:500, Invitrogen cat# A11035). Sections were mounted and coverslipped as above.

#### Mito-RGECO IHC

Sections used to generate figure 4 were immunostained following the direct fluorescence protocol and included the antigen retrieval step prior to permeabilization. Sections were immunostained with rabbit-anti-RFP (1:2000, Rockland cat# 600-401-379, RRID: AB_2209751), chicken-anti-GFP (1:4000, Abcam cat# 13970, RRID: AB_300798), and mouse-anti-RGS14 (1:500, NeuroMab cat# 75-170, RRID: AB_2877352) as a CA2 marker. The secondary antibodies included the following: goat-anti-rabbit (Alexa 546, 1:500, Invitrogen cat# A11035), goat-anti-chicken (Alexa 488, 1:500, Invitrogen cat# A11039) and goat-anti-mouse (Alexa 633, 1:500, Sigma Aldrich cat# SAB4600139). Sections were mounted and coverslipped as above.

### 4.4 mage acquisition

Images used to generate the data in Figs. 1, 2, and 4 were acquired at 16 bit on an inverted Leica Thunder epifluorescent microscope using 10X/0.3 NA, 20X/0.8NA or 63X/1.4NA lenses. Representative images in Figs. 1 and 4 were acquired on a Leica Stellaris SP8 confocal microscope with 63X (1.4NA) lens. For live imaging experiments in Fig. 3, see methods section 4.5.

#### 4.4.1 Mitochondria and dendrites quantification

63× images were acquired at 16 bit and computationally cleared with large volume computational clearing (LVCC) at 92% strength and auto iterations with a limit of 30-40, feature scale of 2689nm, 0.05 regularization, medium optimization, with a refractive index of 1.47 for Prolong Gold. Images from one animal were computationally cleared with instant computational clearing (ICC) for CA1 and CA2, and showed similar trends. As many z slices as needed were acquired to capture as much mitochondria and dendritic segments as feasible. Images were then max projected and quantified in *Fiji*. Secondary and tertiary dendrite segment ROIs were then chosen based on both the TdTomato and MitoTag-GFP channels and cropped, with the primary dendrites excluded. Dendrites with only MitoTag-GFP or only Tdtomato labeling were excluded. All images from one immunolabeling run were scaled to the same fluorescence range in the MitoTag-GFP and the TdTomato channels before converted to 8 bit for thresholding. Otsu auto thresholding was applied to create a MitoTag-GFP binary layer then *Watershed* to separate the objects to best represent the mitochondria in the original image. Image J/Fiji’s *Nucleus Counter* plug-in from Cookbook Microscopy (Collins 2007) was used with a 3×3 median smoothing and a particle range of 10-10,000 to measure mitochondria area. The resulting ROIs were manually edited to best reflect the labeling in the original image. The dendritic segment ROI was outlined using the *Wand Tracing Tool* and manually edited prior to the areas obtained using the *Measure* tool.

Individual mitochondria and dendritic segment areas were normalized to the average of all mitochondria or dendrite areas from within each animal. Total mitochondria area within a dendrite was normalized to the area of that dendritic segment to create a percent occupancy.

#### 4.4.2 OPA1 and MFF quantification

20× computationally cleared images (LVCC, 92% strength, auto iterations, feature scale of 8420nm, 0.05 regularization, and medium optimization) of fixed 40um coronal sections were acquired from naive mice (age 2-4 months) at 16 bit and quantified in Fiji. 11 z slices were max projected and 100 x 100 um ROIs were placed over SR and SLM and 200 x 50 um ROI over SP from 2-4 sections per animal in CA1 and CA2. The fluorescence intensity was obtained using *Measure*. Background signal was subtracted using the negative control section. Sections were then averaged within each animal. Data was normalized to the average of all subregions and layers within each animal.

#### 4.4.3 Mito-RGECO fixed tissue quantification

20x raw (non-computationally cleared) images of CA1 and CA2 subregions of fixed sections from the mitochondrial calcium imaging experiments in 4.2.2 were acquired at 16bit and quantified using Fiji. Images were max projected using all 21 steps, 100um x 100um ROIs were placed over SR and SLM cell layers from 1-5 sections from each animal. The fluorescence intensity was obtained using *Measure*. Each image was background subtracted using the corresponding negative control section of each subregion. SLM fluorescence values were divided by SR values from the same section and subregion to get ratios for CA1 and CA2. Any images showing evidence of cell death or lack of viral expression in all 3 layers were excluded from the quantification.

#### 4.4.4 Mito-RGECO fixed line plot quantification

20x raw (non-computationally cleared) images of CA1 and CA2 subregions were acquired at 16bit and quantified using Fiji. Images were max projected using all 21 steps, three 650um lines were drawn across each subregion per section from SO to SLM, with the middle point of the line consistently placed in the center of SR. The fluorescence intensities across the lines were obtained using *Analyze*, then selecting *Plot Profile*, giving three to four fluorescences intensities per micron for the entire line. The three lines per subregion and per section were averaged to create one line of fluorescence intensities for each subregion and section within animals. Line data was binned by 10um using Prism, and the line fluorescence intensities for each animal were averaged into one line for both subregions. Each animal, for both subregions, was normalized by the average of every point in the line, and then averaged to create one line of fluorescence intensities for CA1 and CA2.

Data was plotted using Prism, and any images showing evidence of cell death or lack of viral expression in all 3 layers were excluded from the quantification.

### 4.5 Electrophysiology: ex vivo brain slice preparation and recording

Experiments were performed on mice aged 6-12 weeks (2-4 weeks post AAV infusion at 4-8 weeks). Cutting and recording solutions were made as described in Helton et al 2019. Mice were deeply anesthetized with 4% isoflurane and transcardially perfused with ice-cold cutting solution containing (in mM): 10 NaCl, 2.5 KCl, 10 D-(+)-glucose, 25 NaHCO_3_, 1.25 NaH_2_PO_4_, 2 sodium pyruvate, 195 sucrose, 7 MgCl_2_, 0.5 CaCl_2_, and saturated with 95% O_2_/5% CO_2_. The brain was rapidly removed and horizontal (CA2) or coronal (CA1) brain slices were cut at 300 µm using a vibratome (VT1000S, Leica). Slices were placed in artificial cerebrospinal fluid (ACSF) containing (in mM): 125 NaCl, 2.5 KCl, 1.25 NaH_2_PO_4_, 25 NaHCO_3_, 20 D-(+)-glucose, 2 Na-pyruvate, 2 MgCl_2_, and 2 CaCl_2_saturated with 95% O_2_/5% CO_2_, and incubated at 33±1°C for 30 min and then at room temperature for > 30 min prior to recording.

For recording, slices were transferred to a submerged recording chamber perfused continuously with 3 ml/min of oxygenated ACSF at 33 ± 1°C. Calcium signals were acquired by widefield fluorescence imaging using an Olympus BX63 microscope equipped with a UM Plan Fluor 10x water immersion objective, 470 nm and 575 nm LEDs (X-Cite Turbo), DAPI/FITC/TRITC/Cy5 Quad filter cube with single band exciters (Chroma), and a Prime 95B CMOS camera (Photometrics). Frames were acquired at 10 s intervals for 10 min including baseline (5 frames) and 110 mM KCl treatment (3 frames) followed by washout with recording aCSF.

#### 4.5.1 Live slice image analysis

Activity-induced calcium signals were quantified as the change in fluorescence relative to baseline (ΔF/F). Baseline (pre) and peak (post) KCl fluorescence intensities were measured with the line plot function in Fiji. The same line was used to measure mito-RGECO and cyto-GCaMP6f per slice. The line was placed so that the middle of the line was squarely in SR, which allowed for general alignment of lines across slices with varying subregion lengths. For the line plots, each line represents one mouse (thin lines) or the average across mice (thick line). For each slice, the pre and post KCl intensity line plot data were normalized to the average pre KCl intensity per channel then binned per 3 µm. Line lengths were different across subregions and slices, therefore distance was normalized by the length of the line. The peak intensities within SP, SR, and SLM were used to quantify ΔF/F per layer and the SLM/SR ratios. If more than one slice per mouse was used, the ΔF/F and SLM/SR values were averaged across slices to generate a single value per mouse.

#### 4.5.3 Post-live imaging tissue processing

Post-recording, brain slices were immediately transferred to 4% paraformaldehyde (PFA) in phosphate buffered saline (PBS) for fixation and incubated at 4°C overnight. Following fixation, slices were washed three times for 5 minutes in 1x PBS to remove residual PFA then either stored in 1x PBS +0.02% azide until processing or blocked overnight at 4°C with gentle agitation in 1x PBS containing 3% normal goat serum (NGS) and 0.3% Triton X100. Blocked slices were transferred to primary antibody solution and incubated for 2 days at 4°C with gentle agitation. Primary antibody solution contained 1x PBS with 3% NGS and 0.3% TritonX-100, rabbit-anti-RFP (1:250, Rockland cat# 600-401-379), chicken-anti-GFP (1:500, Abcam, cat# ab13970), and mouse-anti-RGS14 (1:250, Antibodies Inc, cat# 75-150). Following primary antibody incubation, slices were washed three times for 10 minutes at room temperature (RT) in 1x PBS/0.3% Triton X-100 (PBST) and then transferred to secondary antibody solution to incubate for 48 hours at 4°C or 24 hours at RT, both with gentle agitation. Secondary antibody solution contained 1x PBS with 3% NGS and 0.3% Triton X-100, goat-anti-rabbit Alexa Fluor-conjugated 546 (1:250, Invitrogen cat# A11035), goat-anti-chicken Alexa Fluor-conjugated 488 (1:250, Invitrogen cat# A11039), and goat-anti-mouse Alexa Fluor-conjugated 633 (1:250, Sigma-Aldrich cat# SAB4600139). Slices were then washed three times for 10 minutes in PBST with gentle agitation, then washed once for 10 minutes in 1x PBS + DAPI(1:10,000). Slices were then transferred to 1x PBS to await imaging. 20 minutes prior to image sections were incubated in 60% 2,2’-thiodiethanol (TDE) in a glass bottom 6 well plate to clear the tissue and then imaged on a Leica Thunder with the help of an imaging harp while still submerged in 60% TDE. Thunder processing settings were changed to *Custom* with a refractive index of 1.45 to be able to focus images while in slices in TDE. Slices were imaged immediately after tissue was cleared to avoid deterioration.

## Statistical analyses

All statistical analyses were carried out using Graphpad Prism (v 10.4.1) software, and significance was determined using an alpha level of 0.05.

## DECLARATIONS

### Authors’ contributions

Conceptualization, SF, SAS; Methodology, KP, LT, MA, RT, SF, SAS; Formal Analysis, LT, MA, RT, SF, SAS; Investigation, KP, LT, MA, RD, RT, SF, SAS; Writing – Original Draft, LT, MA, SF, Writing – Review & Editing, KP, LT, MA, RD, RT, SF, SAS; Visualization, LT, MA, RT, SF, SAS; Supervision, SF, SAS; Funding Acquisition, SF, SAS.

## Acknowledgements

The authors acknowledge resources and support from the Virginia Tech animal care staff and Cellular and Molecular Imaging Core, part of the Fralin Biomedical Research Institute at VTC. This research would not be possible without the use of the Leica Stellaris SP8 Microscope.

